# SparseAEH: Scalable autoregressive expression histology discovery in spatial transcriptomics via sparse Gaussian kernels

**DOI:** 10.64898/2026.01.22.701193

**Authors:** Tianjie Wang, Lingyu Li, Yuanhua Huang

**Affiliations:** School of Biomedical Sciences, The University of Hong Kong, Hong Kong SAR; Department of Statistics and Actuarial Science, The University of Hong Kong, Hong Kong SAR

## Abstract

**Summary:** Spatial clustering of genes is an effective way to elucidate histological or spatial patterns of marker expressions or ligand-receptor interactions. However, its computational complexity hinders its application to the advanced spatial transcriptomics (ST) technologies with high resolution and throughput. Here, we present SparseAEH, a highly scalable algorithm and implementation of the Gaussian process mixture model, by leveraging block-wise sparse covariance for approximating the likelihood. This acceleration enables the analysis of dense ST datasets within practical time constraints while maintaining accuracy. Benchmarking on both simulated and experimental data demonstrates that our method achieves significantly less computational time with comparable accuracy to previous methods, highlighting its potential as a powerful tool for the exploration of high-resolution ST data.

**Availability and implementation:** SparseAEH is implemented in Python and is publicly available at https://github.com/jackywangtj66/SparseAEH.

## 1 Introduction

Spatial transcriptomics (ST) has become a major tool for understanding tissue biology by *in situ* profiling the gene expression with sequencing or imaging readouts (Ståhl *et al*., 2016; Vickovic *et al*., 2019). These ST technologies are undergoing rapid development, increasing the resolution of the profiling bin and the size of the profiling area, resulting in tens to hundreds of thousands of spatial spots, e.g., by sequencing like VisiumHD (Oliveira *et al*., 2025), by imaging like Xenium (Janesick *et al*., 2023), or by histology-based imputation like our FineST (Li *et al*., 2026). While such high-resolution data enables more fine-grained analysis for a better understanding of intercellular interactions, the large data size introduces a major computational burden for spatial pattern discovery. For deep learning-based models, this computational bottleneck can be partly alleviated by using more advanced GPU hardware with smaller batches, while for statistical models, mathematical approximation can be a surprisingly well-balanced solution. One example is detecting spatial correlation, where SpatialDM using *z*-score approximation achieves over a thousand times speedup and enables analysis of a million spots (Li *et al*., 2023).

Identifying spatial expression patterns underlying ST data is essential for identifying tissue domains and understanding how gene regulation varies across different microenvironments. Besides conventional latent variable methods, multiple strategies have been proposed to incorporate spatial dependency, either deterministically (e.g., using a graph neural network, either in the encoder or decoder) or probabilistically often via Gaussian Process (GP) models (Rasmussen and Williams, 2005). Specifically, GPs are powerful and interpretable models, encoding these spatial-dependent relationships within their covariance matrices. Spatial pattern discovery with GPs is often done in two orthogonal ways. One is the GP latent variable model, a type of factor analysis model, treating cells as samples, like MEFISTO (Velten *et al*., 2022) and NSF (Townes and Engelhardt, 2023), and consequently returning factors with spatial information. The other is the GP mixture model, using genes as samples, like SpatialDE (Svensson *et al*., 2018), to perform clustering over a set of (spatially variable) genes. Nevertheless, despite their enhanced scalability compared to earlier techniques (e.g., with variational inference in SpatialDE), these methods still fundamentally require operation on the large covariance matrices, hindering them from analysing increasingly high-dimensional data effectively.

To address this challenge and with a focus on the GP mixture model, we here propose a highly efficient algorithm, SparseAEH, to perform clustering over features (e.g., genes). While building upon a common Gaussian prior assumption and employing the traditional Expectation-Maximization (EM) (Dempster *et al*., 1977) algorithm, our method introduces a novel strategy for approximating probabilities by leveraging spatial information, in a similar spirit to sparse GP with inducing points. This approach effectively circumvents the direct computation of large covariance matrices. Our experiments on both simulated and real-world datasets demonstrate SparseAEH’s superior accuracy and computational efficiency, highlighting its great potential for handling increasingly high-dimensional ST data.

## 2 Results

### 2.1 Efficient probability estimation

We first present the schematic workflow of SparseAEH, which involves the initial clustering of spots, aggregation into superspots based on spatial proximity, and subsequent sparsification of the covariance matrix by retaining only spatially relevant entries (**Fig**. 1A). To assess the accuracy and computational efficiency of SparseAEH in approximating the likelihood of high-dimensional data, we used synthetic high-dimensional Gaussian data (generated from a kernel built on 5,000 spatial locations), and estimated likelihoods with both the sparse covariance used in SparseAEH and the raw full covariance by Scipy’s built-in function as a reference (same as used in SpatialDE). As shown in **Fig**. 1B, SparseAEH demonstrates highly accurate approximation of the reference likelihoods. Furthermore, we investigated the computational scalability of both methods by varying the data dimension. As shown in **Fig**. 1C, SparseAEH exhibited superior scaling performance, particularly when the dimension exceeded 2,000, showcasing its ability to handle high-dimensional data efficiently.

**Fig. 1.**
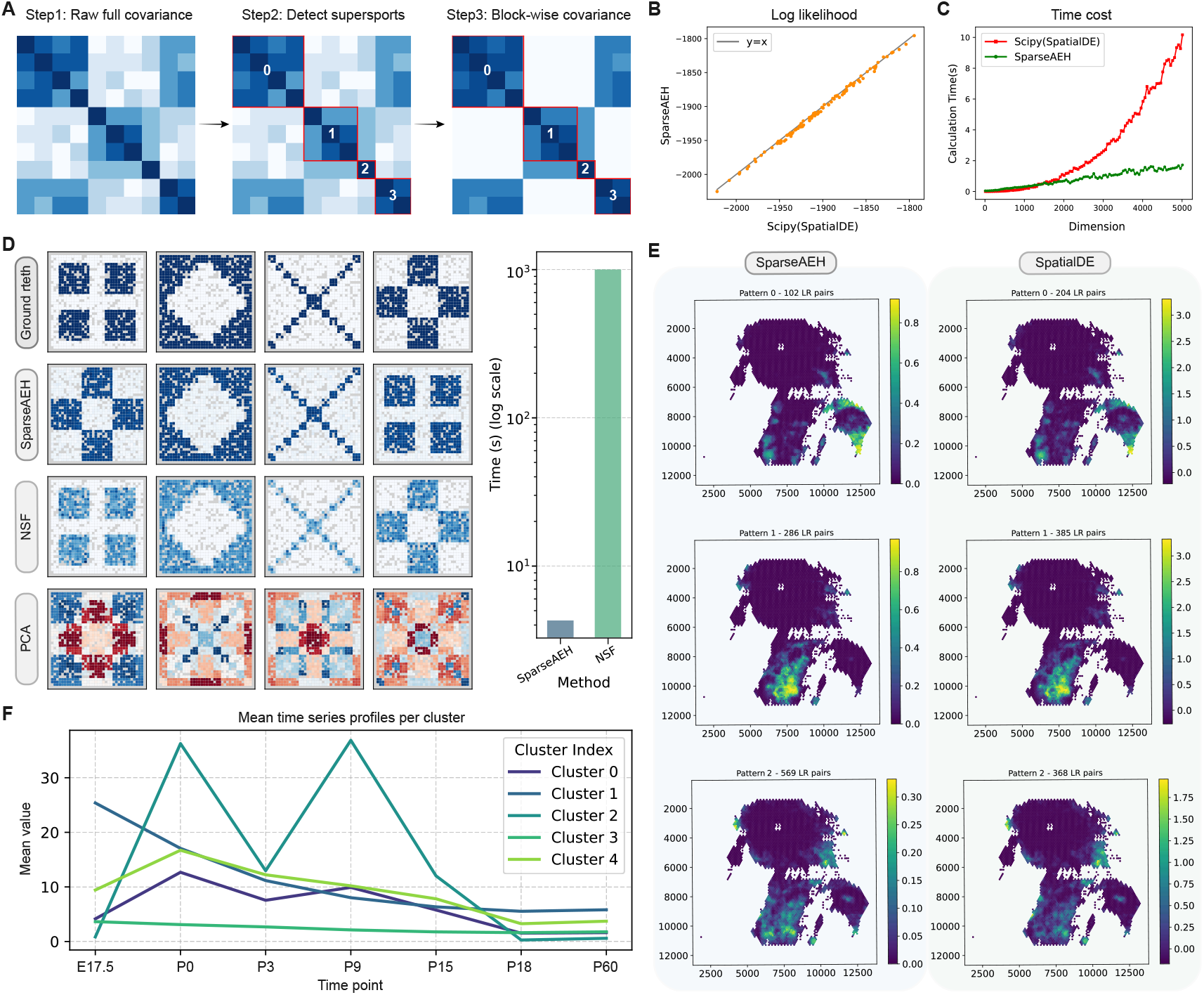
Overview and benchmarking of SparseAEH across diverse clustering tasks, including simulated data, a real-world 10x Visium NPC dataset, and an embryonic beta cell single-cell temporal dataset. (A) Typical procedure of SparseAEH, during which the spots are first roughly divided into clusters. Step 1: Original full covariance matrix. Step 2: Spots are grouped into superspots based on spatial proximity. Step 3: Keep entries only between the identified spatially related superspots and ignore the others. (B) Accurate log-likelihoods approximation on simulated data by SparseAEH, compared to Scipy (with raw full covariance, same as SpatialDE), showing close alignment with the *y* = *x* line. (C) Comparison of running time on simulated data up to 5,000 dimensions between SparseAEH and its counterpart with full covariance (by Scipy, same as SpatialDE). When the dimension exceeds 2,000, SparseAEH begins to show a significant advantage. (D) Performance of pattern detection among various methods on simulated data. The top 4 patterns are the ground truth. SparseAEH and NSF can both identify the ground truth. However, SparseAEH only spent 4.3 seconds, while NSF needed 1004.1 seconds. (E) The clustering results of SparseAEH and SpatialDE on the 4*×* super-resolved NPC dataset from FineST (with 5,039 spots and 957 LR pairs), using respectively 12.253 seconds and 21,654.154 seconds. The detected cluster and spatial patterns were visually similar. (F) The results of SparseAEH on a time-series data on the embryo beta cells, showing its applicability to temporal data.

### 2.2 Performance on simulated dataset

To evaluate SparseAEH’s performance in recovering spatial patterns, we conducted experiments using a simulated dataset, following the pipeline outlined in Townes and Engelhardt (2023). **Fig**. 1D presents the clustering results of different methods on the ‘quilt’ dataset, consisting of four overlapping spatial patterns. Notably, SparseAEH accurately reconstructed all four patterns with significantly less computational time than NSF (4.3 seconds by SparseAEH vs 1,004.1 seconds by NSF; over 200 speedups), while MEFISTO failed to distinguish the true patterns apart.

### 2.3 Application to real-world ST dataset

We also applied SparseAEH to a 10x Visium nasopharyngeal carcinoma (NPC) (Gong *et al*., 2023) ST dataset consisting of 1,331 spots. First, the data was pre-processed with FineST to enhance resolution to the sub-spot level, which consequently increased its dimension to around 5,000. Subsequently, SpatialDM was employed to discover spatially interacting ligand-receptor (LR) pairs, yielding 931 spatially significant LR pairs. These LR pairs then served as input for SparseAEH to uncover underlying cell-cell communication patterns. Using SpatialDE as a benchmark, SparseAEH produced results comparable to SpatialDE, yet with significantly lower computational time (**Fig**. 1E).

### 2.4 Capability of temporal dataset analysis

Besides spatial dependency, the GP model is also a common choice for temporal modelling, e.g., for splicing via time-series RNA-Seq (Huang and Sanguinetti, 2016). Here, we showcase that, as a side product, SparseAEH is readily applicable for temporal datasets, where we only need to duplicate the time variable into a 2-dimensional input format to align with the algorithm’s structure. To demonstrate this, we evaluated SparseAEH on a pancreatic beta cell dataset (Qiu *et al*., 2017), comprising 575 cells measured at 7 developmental time points from embryonic 17.5 day (E17.5) to postnatal 60 day (P60). As illustrated in **Fig**. 1F, SparseAEH identifies distinct clusters of cells with characteristic temporal expression profiles, capturing dynamic changes in gene expression over time.

## 3 Methods

### 3.1 Gaussian mixture model with spatially-derived full covariance

Let **y** = (*y*_1_, *y*_2_, …, *y*_*N*_) correspond to a vector of gene expression over *N* spots **X** = (**x**_1_, **x**_2_, …, **x**_*N*_), where each spot **x**_*i*_ is represented by 2-dimensional vector (*x*_*i*1_, *x*_*i*2_). The expression matrix **Y** consists of G expression vectors (**y**_1_, **y**_2_, …, **y**_*G*_). Assume that there are *K* underlying patterns for the Gaussian mixture models, and the probability of each expression vector being sampled from the *k*-th pattern is *π*_*k*_, such that 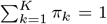. A binary matrix **Z** indicates the gene *g* is explained by pattern *k* if *z*_*g,k*_ = 1. The expression model can be formulated as the following Gaussian mixture model:

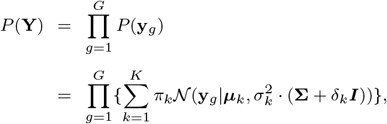

where ***µ***_*k*_ denotes the mean vector of the *k*-th Gaussian pattern and **Σ** denotes the covariance matrix universal for all patterns, realized by the RBF kernel:

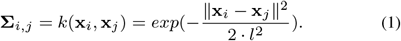

We propose the SparseAEH algorithm as follows (see **Algorithm 1**):

#### Algorithm 1

SparseAEH Initialization

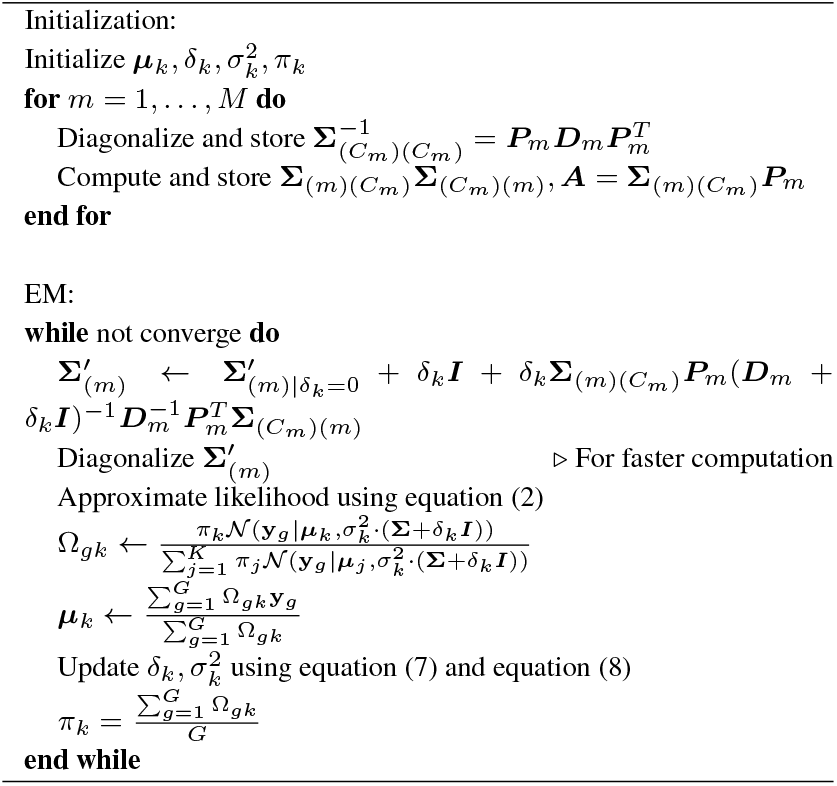

### 3.2 Sparse approximation with block-wise covariance

Partition the *N* spots into *M* sets of superspots, {*S*_1_, *S*_2_, …, *S*_*M*_ } such that every spot belongs to one of the superspots. Let {**X**_*i*_} be the set of spots belonging to {*S*_*i*_} and

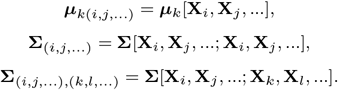

Let **y**_*gi*_ be the sub-vector of **y**_*g*_ corresponding to the spots {*S*_*i*_}. To limit the number of other spots that each spot is correlated with, let *C*_*i*_ be the set of indices of superspots that *S*_*i*_ is dependent on. Without loss of generality, we can further assume that ∀*i* ∈ *C*_*j*_, *i < j*.

Let’s define the conditional mean and covariance terms to simplify the main expression:

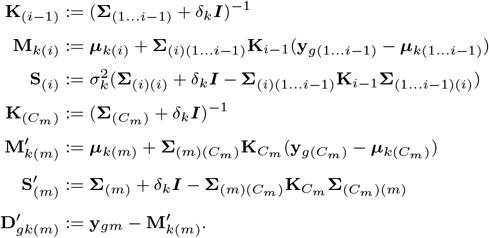

Then the full expression can be written as:

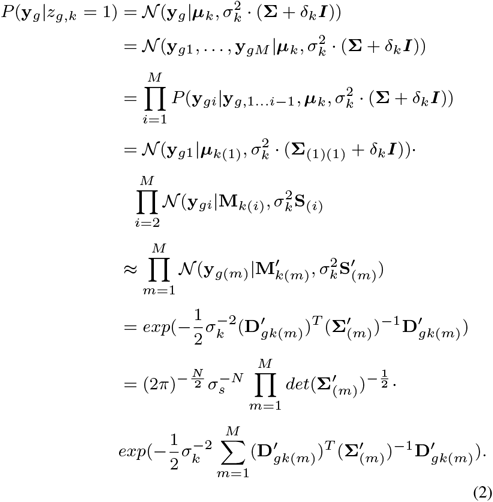

### 3.3 Estimation with EM algorithm

With the block-wise covariance approximation we stated in Section 2, we are ready to apply the EM algorithm to solve the clustering problem, in which the major difficulty for the high-dimensional case lies in estimating the posterior:

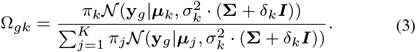

Take derivative with respect to mean yields the local maximum likelihood estimation (MLE) of ***µ***_*k*_:

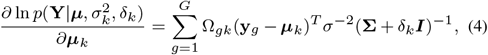

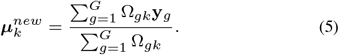

To update δ_*k*_ and 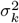, we minimize the Frobenius norm:

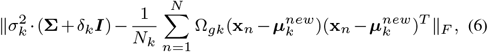

Where 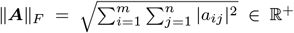 for any matrix ***A*** ∈ K^*m×n*^ (a field of either real or complex numbers). The optimal solution can also be calculated straightforwardly:

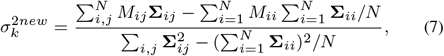

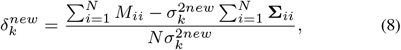

where 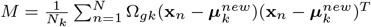

To facilitate easier updates in the algorithm, we also derived:

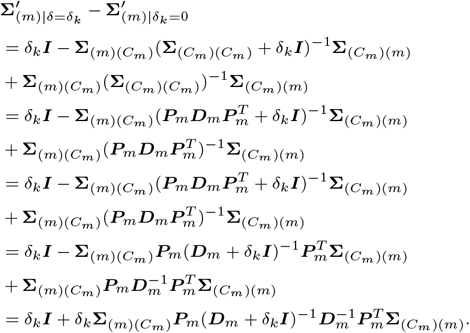

## 4 Conclusion

In this work, we present SparseAEH, an EM-algorithm-based method for spatial pattern identification. We discovered that ignoring the impact from distant spatial locations maintains significant accuracy while substantially reducing computational time. We applied SparseAEH on simulated and real datasets, confirming its efficacy in uncovering intricate spatial patterns. Furthermore, SparseAEH also demonstrates applicability to temporal data analysis.

Limited by the basic assumption of a mixture model, SparseAEH can not be used to perform dimension reduction. While our approximation logic significantly enhances the efficiency of probability computation, it does not readily extend to direct operations on the full covariance matrix, such as calculating the matrix inverse. In the future, we expect to derive more methods to manipulate the covariance matrix based on a similar idea.

Besides, although SparseAEH can deal with both spatial and temporal data, just like previous methods, it still lacks the ability to analyze ST data from different time points collectively. Identifying genes that share similar spatio-temporal patterns can offer great help in tracking the cell fate, identifying dynamic cell-cell communication, and elucidating dynamic biological pathways. With its strong scalability, SparseAEH can serve as a good foundation for future development of comprehensive methods.

## Acknowledgements

We would like to thank Dr. Yijun Liu and Dr. Ruiyan Hou for their support in providing and assisting us in understanding the temporal dataset.

## Author contributions

Tianjie Wang (Data curation [equal], Methodology [equal], Formal analysis [equal], Software [Lead], Visualization [equal], Validation [equal], Writing—review & editing [equal]), Lingyu Li (Data curation [equal], Visualization [equal], Validation [equal], Writing—review & editing [equal]), and Yuanhua Huang (Conceptualization [lead], Investigation [lead], Methodology [equal], Formal analysis [equal], Validation [equal], Funding acquisition [lead], Project administration [lead], Supervision [lead], Writing—review & editing [equal]) *Conflict of Interest:* none declared.

## Data availability

All data underlying this resource, as well as the code used for its assembly, are publicly available at https://github.com/jackywangtj66/SparseAEH.

